# Rescuable sleep and synaptogenesis phenotypes in a *Drosophila* model of O-GlcNAc transferase intellectual disability

**DOI:** 10.1101/2023.06.28.546900

**Authors:** Ignacy Czajewski, Laurin McDowall, Andrew Ferenbach, Daan M. F. van Aalten

**Affiliations:** Division of Molecular, Cell and Developmental Biology, School of Life Sciences, University of Dundee, Dundee, UK; Department of Molecular Biology and Genetics, Aarhus University, Aarhus, DK

## Abstract

O-GlcNAcylation is an essential intracellular protein modification mediated by O-GlcNAc transferase (OGT) and O-GlcNAcase (OGA). Recently, missense mutations in *OGT* have been linked to intellectual disability, indicating that this modification is important for the development and functioning of the nervous system. However, the processes that are most sensitive to perturbations in O-GlcNAcylation remain to be identified. Here, we uncover quantifiable phenotypes in the fruit fly *Drosophila melanogaster* carrying a patient-derived OGT mutation in the catalytic domain. Hypo-O-GlcNAcylation leads to defects in synaptogenesis and reduced sleep stability. Both these phenotypes can be partially rescued by genetically or chemically targeting OGA, suggesting that a balance of OGT/OGA activity is required for normal neuronal development and function.

## Introduction

Intellectual disability (ID) is a disorder affecting around 1% of the population globally^1^, characterised by an intelligence quotient lower than 70 accompanied by reduced adaptive behaviour^2^. Recently, mutations in the X chromosome gene *OGT* were identified as causal for ID, a condition termed OGT congenital disorder of glycosylation (OGT-CDG)^3–7^. Patients with OGT-CDG present with diverse signs of varying penetrance, such as microcephaly and white matter abnormalities, as well as non-neurological signs such as clinodactyly, facial dysmorphism, and developmental delay, manifesting as low birth weight and short stature^4^. Beyond ID, non-morphological signs of pathogenic *OGT* mutations include behavioural problems as well as sleep abnormalities and epilepsy^4, 8^.

*OGT* encodes a nucleocytoplasmic glycosyltransferase, O-linked β-N-acetyl glucosamine (O-GlcNAc) transferase (OGT), a multifunctional protein composed of two domains: a tetratricopeptide repeat (TPR) domain and a catalytic domain^9–11^. Mutations affecting either domain have been identified in patients with OGT-CDG, though clinical manifestation of the disorder does not appear to segregate with the domain affected^4^, suggesting a common disease mechanism. The N-terminal TPR domain is believed to confer substrate specificity for the glycosyltransferase function of OGT^12–14^ and is important for non-catalytic functions of the protein^15, 16^. The catalytic domain fulfils two known functions, the transfer of O-GlcNAc onto serine and threonine residues of nucleocytoplasmic proteins (O-GlcNAcylation)^17, 18^, and the proteolytic activation of Host Cell Factor 1 (HCF-1)^19, 20^, a known ID-associated protein^21^. While the latter function of OGT potentially contributes to the pathogenicity of some *OGT* mutations^5^, not all patient mutations have been found to affect HCF-1 processing, neither *in vitro* nor when modelled in stem cells^7, 8, 22^. Overall, the role of altered O-GlcNAcylation in OGT-CDG pathogenicity remains an open question, as many of the other functions fulfilled by this protein have the potential to contribute to ID.

O-GlcNAcylation is a dynamic modification occurring on around 5000 proteins in the human proteome^23^. The dynamic nature of the modification is conferred by O-GlcNAcase (OGA), which opposes OGT, catalysing the removal of O-GlcNAc^24, 25^. O-GlcNAcylation has been extensively implicated in neuronal development, functioning and disease^26–31^ and is therefore likely to play a key role in the pathogenicity of OGT-CDG. The first evidence for the requirement for OGT in development was the study of *Drosophila melanogaster OGT*, *super sex combs* (*sxc*), as a Polycomb group (PcG) gene, amorphic mutations of which were found to result in defects in body segment determination^32^, a function later ascribed to its glycosyltransferase activity^33^. The role of O-GlcNAcylation in PcG function is known to be important for normal neuronal development, and highly sensitive to perturbations. For example, maternal hyperglycaemia can drive increased O-GlcNAcylation in the embryo altering neuronal maturation and differentiation patterns through altered PcG function^34^. Multiple additional core developmental regulators have been found to require O-GlcNAcylation for appropriate function, with deregulation of the modification affecting stem cell maintenance through core pluripotency factors such as Sox2^35–37^, cell fate determination through STAT3^38^ and Notch signalling^30^ and neuronal morphogenesis through the protein kinase A signalling cascade^39^. Additionally, O-GlcNAcylation is known to play an important role in neuronal functioning related to memory formation^40, 41^. For example, elevating O-GlcNAcylation in sleep deprived zebrafish or mice can reverse memory defects associated with a lack of sleep^42, 43^. The extent of the role of OGT in memory formation is not fully understood, although several proteins important for this process are modulated by O-GlcNAcylation, such as CREB^44^ or CRMP2^40^. Therefore, a key unanswered question regarding the aetiology of OGT-CDG is the contribution of the developmental roles of OGT relative to its role in the functioning of the adult nervous system.

With the large number of functionally O-GlcNAcylated proteins and thousands more which remain uncharacterised, identifying the most important processes controlled by O-GlcNAcylation remains challenging. Patient mutations in the catalytic domain present a unique opportunity to better understand processes most sensitive to defective O-GlcNAc cycling. Therefore, we set out to model catalytic domain intellectual disability mutations in *Drosophila melanogaster* and characterise their phenotypic effect. *Drosophila* OGT (*Dm*OGT) is highly similar to its human ortholog, with 73% amino acid identity and a high degree of structural similarity^45^. However, in the fly OGT does not catalyse HCF-1 proteolytic activation, a function fulfilled instead by taspase 1^46^, eliminating this function of OGT as a confounding variable in understanding the role of O-GlcNAcylation in ID. Previous work modelling OGT-CDG mutations in *Drosophila* has demonstrated that OGT-CDG catalytic domain mutations can reduce global O-GlcNAcylation in adult tissue^5^, which is linked with defects in habituation and synaptogenesis^47^. Here, we demonstrate that a recently discovered ID associated catalytic domain mutation in *OGT* (resulting in the amino acid substitution C921Y^22^) can reduce O-GlcNAcylation throughout development in *Drosophila*, which can be rescued by genetically or pharmacologically abolishing or reducing OGA activity, respectively. We find a strong effect of *sxc* mutations on larval neuromuscular junction (NMJ) development, which can be partially reversed by inhibiting or abolishing OGA catalytic activity. Additionally, we demonstrate that a catalytic domain mutation in *sxc* can negatively impact sleep, reducing sleep bout duration. This phenotype can be rescued by abolishing OGA activity and partially reversed by inhibiting OGA in adulthood, suggesting that some aspects of OGT-CDG may not be developmental in origin.

## Results

### An OGT-CDG mutation reduces global O-GlcNAcylation throughout Drosophila development

To investigate the contribution of reduced O-GlcNAcylation to phenotypes relevant to OGT-CDG, catalytic domain mutations found in patients were modelled in *Drosophila* using CRISPR-Cas9 mutagenesis. The previously published *sxc^N595K^* (equivalent to human N567K)^5^ and the newly generated *sxc^C941Y^* (equivalent to human C921Y) mutant strains were used to assay the effects of OGT-CDG mutations on global O-GlcNAcylation in adult flies. Consistent with previous reports, O-GlcNAcylation in lysates from adult heads was found to be significantly reduced in the *sxc^N595K^* mutant compared to a control genotype (Fig. 1A)^5^. The newly generated *sxc*^*C941Y*^ mutant strain presented with a significantly more severe reduction in global O-GlcNAcylation, to roughly 40% of the control genotype. This reduction in O-GlcNAcylation was observed despite a modest, yet significant, increase in OGT protein relative to the control genotype. As the reduction in O-GlcNAcylation was modest in the *sxc*^*N595K*^ line, a previously generated catalytically dead mutant strain (*sxc^K872M^*) was further characterised alongside the newly generated *sxc*^*C941Y*^ variant^48^, to control for allele specific effects. The *sxc^K872M^* genotype was previously found to be recessive lethal at the late pupal stages^48^, therefore, for this genotype O-GlcNAcylation and OGT levels were only assayed at embryonic and larval stages. Both *sxc*^*C941Y*^ and *sxc^K872M^* stage 16-17 embryos present with significantly reduced O-GlcNAcylation and increased OGT (Fig. 1B). As *sxc^K872M^* embryos were derived from heterozygous parents, O-GlcNAcylation seen in these embryos is likely largely due to maternally contributed wildtype *sxc* gene product^32, 49^. By the third instar larval stage of development, the difference in O-GlcNAcylation between the *sxc*^*C941Y*^ and *sxc^K872M^* genotypes is more pronounced. *Sxc^K872M^* larvae present with significantly lower O-GlcNAcylation than both the control and *sxc*^*C941Y*^ genotype (Fig. 1C). O-GlcNAcylation in the *sxc*^*C941Y*^ larvae remains significantly reduced relative to the control genotype, as at all other stages of development assayed. Surprisingly, at this stage of development, *sxc*^*C941Y*^ larvae do not present with significantly elevated *Dm*OGT protein levels. Strikingly, the mean *Dm*OGT protein levels in *sxc^K872M^* larvae are over 8 times higher than in the control genotype.

**Figure 1:**
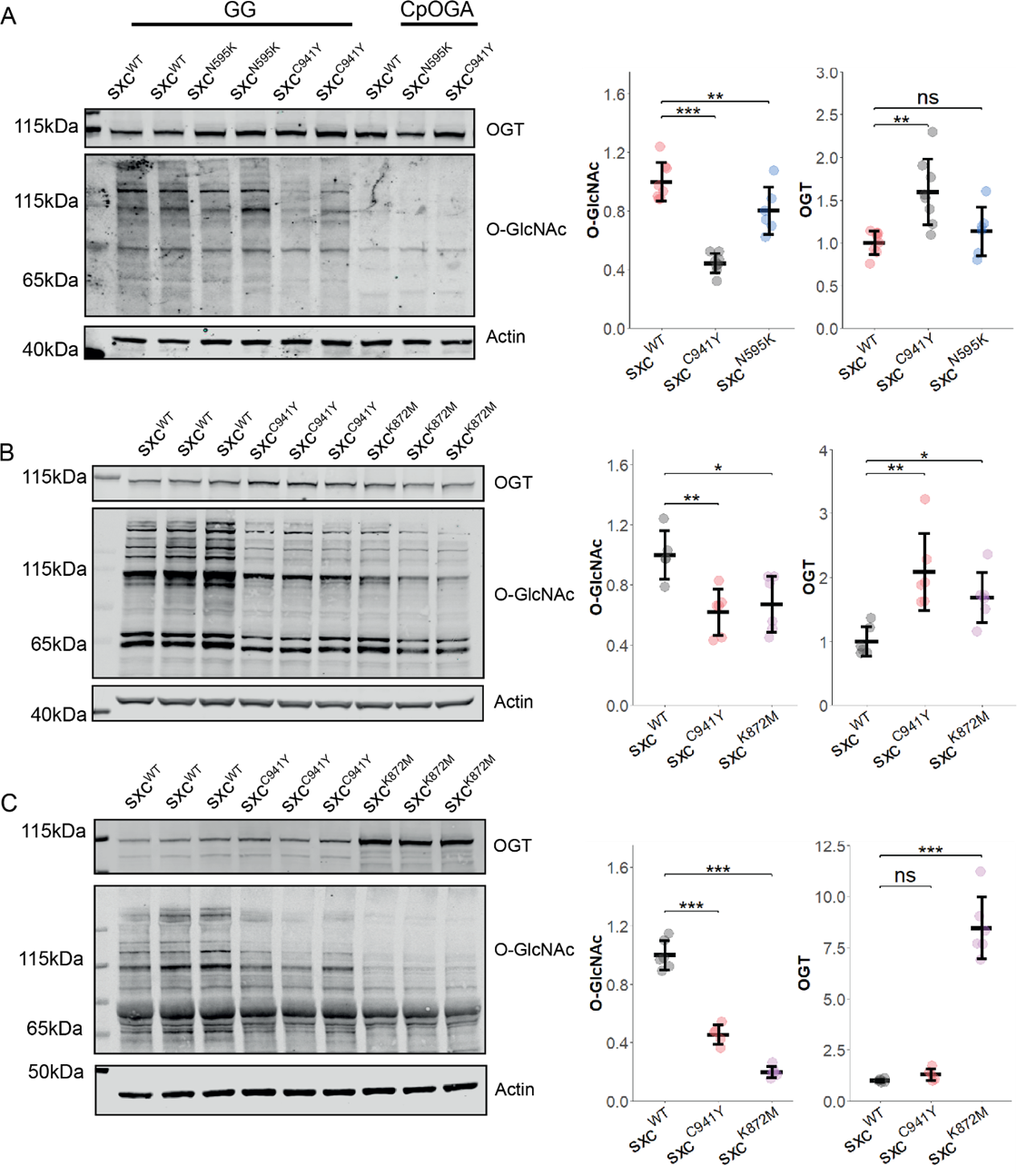
**A**) Representative Western blot of *sxc^WT^* (n = 8), *sxc*^*C941Y*^ (n = 8), and *sxc*^*N595K*^ (n = 6) adult head lysates and quantification (mean ± standard deviation) of OGT and O-GlcNAcylation immunoreactivity normalised to the both the loading and genotype control. *Clostridium perfringens OGA* (CpOGA) treated lanes demonstrate the specificity of the O-GlcNAc antibody (RL2) used when compared to lysates treated with the OGA inhibitor GlcNAc statin G (GG). A significant intergroup difference in O-GlcNAcylation was observed (F(2,19)= 42.82, *p* < 0.001), with post hoc analysis revealing a significant reduction in O-GlcNAcylation in both *sxc*^*N595K*^ (p_adj_ < 0.05) and *sxc*^*C941Y*^ (*p_adj_* < 0.001) flies relative to the control genotype, and a significant difference between the mutant strains (*p_adj_* < 0.001). A significant intergroup difference was also observed for OGT levels (F(2,19) = 9.137, *p* < 0.01), however, post hoc analysis revealed this was only due to a significant increase in OGT in *sxc*^*C941Y*^ flies (*p_adj_*< 0.01) **B)** Representative Western blot of *sxc^WT^* (n = 6), *sxc*^*C941Y*^ (n = 6), and *sxc^K872M^* (n = 6) lysates from stage 16-17 embryos along with OGT and O-GlcNAc quantification. A significant decrease in O-GlcNAcylation (F(2,14) = 8.014, *p* < 0.01) was observed for both *sxc*^*C941Y*^ (*p_adj_* < 0.01) and *sxc^K872M^* (*p_adj_* < 0.05) embryos, accompanied by a significant increase in OGT (F(2,14) = 9.49, *p* < 0.01) for both genotype (*p_adj_* < 0.01 and *p_adj_*< 0.05, respectively). **C)** Representative Western blot and quantification of lysates from *sxc^WT^* (n = 6), *sxc*^*C941Y*^ (n = 5), and *sxc^K872M^* (n = 6) third instar larvae, demonstrating a significant decrease in O-GlcNAcylation for both *sxc*^*C941Y*^ and *sxc^K872M^* larvae (F(2,14) = 184.5 *p* < 0.001, *p_adj_* < 0.001 and *p_adj_* < 0.001, respectively) and a decrease in OGT in *sxc^K872M^* larvae (F(2,14) = 122.6, *p* < 0.001, *p_adj_* < 0.001). * p < 0.05, ** p < 0.01, *** p < 0.001.

To determine whether the phenotypic consequences of loss of O-GlcNAc transferase function in the *sxc*^*C941Y*^ mutant flies results in similar phenotypic consequences as a previously characterised *Drosophila* line carrying a hypomorphic mutation in sxc (*sxc^H537A^*), flies were assayed for scutellar bristle development^48^. *sxc*^*C941Y*^ flies were found to also present with an increased penetrance of ectopic bristles on the scutellum, with 31% of *sxc*^*C941Y*^ flies presenting with one or more additional bristles, while in the control genotype this only occurred in 8% of flies (Fig. S1A). Taken together, these results demonstrate that hypo-GlcNAcylation due to OGT-CDG variants can be modelled in *Drosophila*. Further supporting the hypothesis that reduced OGT catalytic activity is causal in phenotypes seen in ID, a patient mutation modelled in *Drosophila* results in a similar phenotype as rational mutagenesis of a key *Dm*OGT catalytic residue.

### Pharmacological rescue of O-GlcNAc levels in sxc^C941Y^ flies

To evaluate whether reduced O-GlcNAcylation in *sxc* mutants with impaired catalytic activity can be rescued to control levels, we sought to elevate O-GlcNAcylation through both genetic and pharmacological means. First, to demonstrate that O-GlcNAcylation can be rescued in flies with impaired *Dm*OGT catalytic activity by abolishing OGA activity, *sxc* mutant flies were crossed with an *Oga* knockout strain (*Oga^KO^*)^50^. When assayed by Western blot, we found that knocking out OGA led to a marked increase in O-GlcNAcylation in lysates from adult heads in the *sxc*^*C941Y*^ line, above levels seen in the control genotype (Fig. 2A). To assay whether this rescue of O-GlcNAcylation could reverse a phenotype caused by reduced O-GlcNAc transferase activity, we compared the number of scutellar bristles in *sxc^WT^*, *sxc*^*C941Y*^, *sxc*^*C941Y*^;*Oga^KO^*, and *Oga^KO^* flies. Surprisingly, we found that despite the *Oga^KO^* allele having no effect on its own, *sxc*^*C941Y*^;*Oga^KO^* flies had an increased penetrance of ectopic scutellar bristles beyond what we observed for *sxc*^*C941Y*^ flies (Fig. S1B).

**Figure 2:**
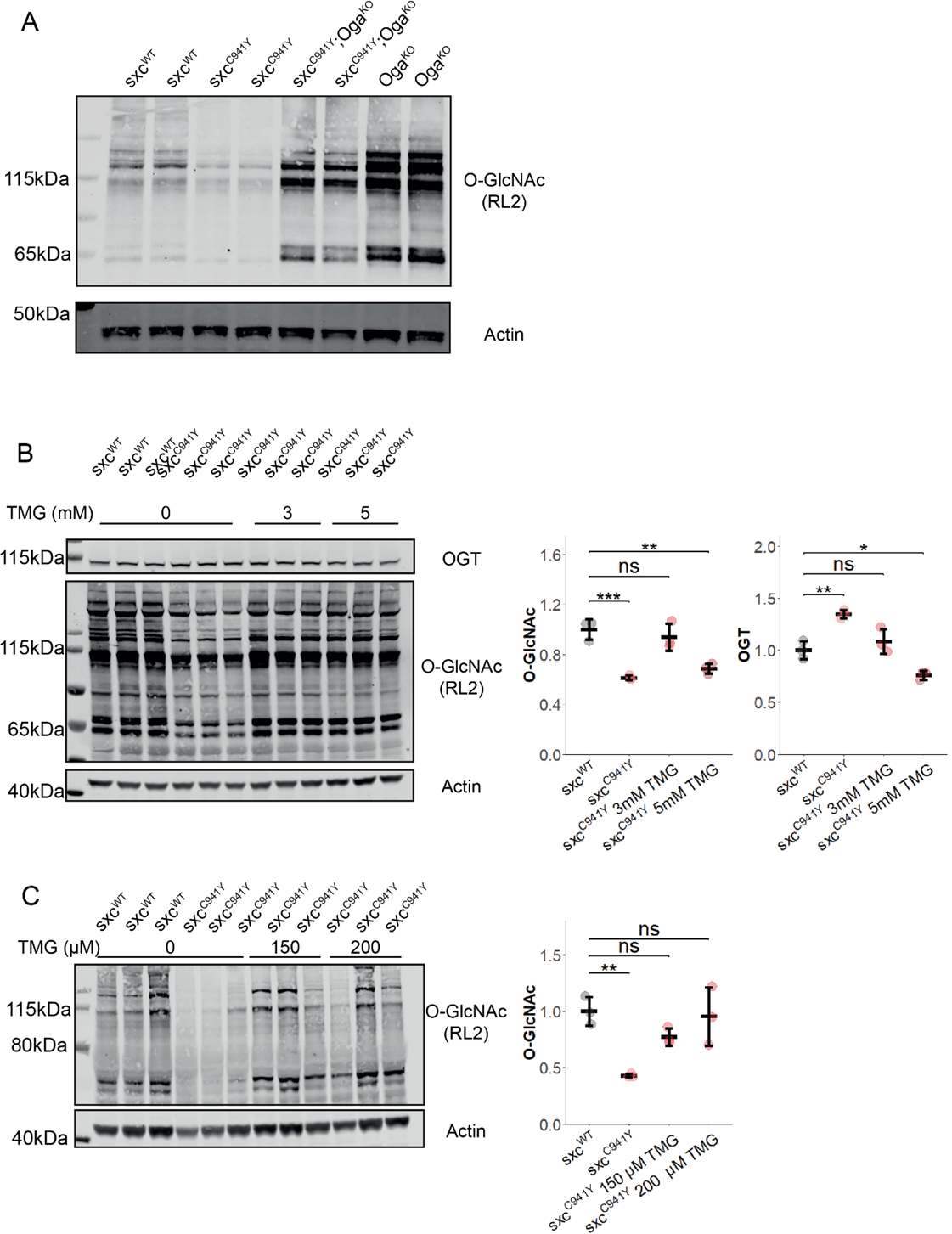
**A**) Representative Western blot (of three) of *sxc^WT^*, *sxc*^*C941Y*^, *sxc*^*C941Y*^;*Oga*^*KO*^ and *Oga^KO^* adult head lysates, immunolabelled with RL2 to detect O-GlcNAcylation and actin as a loading control. **B**) Western blot and quantification of adult head lysates of *sxc^WT^*, *sxc*^*C941Y*^ vehicle, *sxc*^*C941Y*^ fed 3 mM Thiamet G (TMG) and *sxc*^*C941Y*^ fed 5 mM TMG (n = 3), immunolabelled for O-GlcNAcylation using the RL2 antibody, OGT, and actin as a loading control. A significant intergroup difference was observed for both O-GlcNAcylation (F(3,8) = 20.86, *p* < 0.001) and OGT (F(3,8) = 27.28, *p* < 0.001) levels, with post hoc analysis revealing that O-GlcNAcylation and OGT levels were not significantly different between *sxc^WT^* flies and *sxc*^*C941Y*^ flies fed 3 mM TMG (*p_adj_* = 0.75 and *p_adj_*= 0.57, respectively). Both *sxc*^*C941Y*^ flies fed a vehicle control and 5 mM TMG present with significantly different O-GlcNAcylation (*p_adj_* < 0.001 and *p_adj_* < 0.01, respectively) and OGT (*p_adj_* < 0.01 and *p_adj_* < 0.05, respectively) levels. **C**) Western blot and quantification of third instar larval lysates of *sxc^WT^*, *sxc*^*C941Y*^ vehicle, *sxc*^*C941Y*^ fed 150 μM TMG and *sxc*^*C941Y*^ fed 200 μM TMG (n = 3), immunolabelled for O-GlcNAcylation using the RL2 antibody and actin as a loading control. O-GlcNAcylation significantly differed between groups (F(3,8) = 9.11, *p* < 0.01), with both 150 μM and 200 μM TMG rescuing O-GlcNAcylation levels in *sxc*^*C941Y*^ larvae to be no longer significantly different relative to the control genotype (*p_adj_* =0.31 and *p_adj_* =0.98, respectively) and 200 μM TMG treatment significantly elevating O-GlcNAcylation relative to the untreated *sxc*^*C941Y*^ larvae (*p_adj_* < 0.05). * p < 0.05, ** p < 0.01, *** p < 0.001.

We next set out to identify concentrations at which the OGA inhibitor Thiamet G (TMG) ^51^ would restore *sxc*^*C941Y*^ global O-GlcNAcylation to control levels. To elevate O-GlcNAcylation in adult *Drosophila,* young adult flies were placed on food supplemented with TMG for 72 h prior to analysis by Western blotting (Fig. 2B). After assaying varying concentrations of OGA inhibitor, we found that global O-GlcNAcylation was rescued to control levels in *sxc*^*C941Y*^ flies fed 3 mM TMG for 72 h. Paradoxically, a higher 5 mM concentration did not have the same effect. Flies fed this higher concentration of TMG were found to have significantly decreased global O-GlcNAcylation relative to the control genotype, though this appeared to be due to an alteration in the pattern of O-GlcNAcylation with some substrates retaining elevated O-GlcNAcylation relative to *sxc*^*C941Y*^ flies fed standard food (Fig. S2A). Accompanying elevated O-GlcNAcylation, TMG treatment resulted in decreased levels of *Dm*OGT. For *sxc*^*C941Y*^ flies fed 3 mM TMG, *Dm*OGT protein levels were rescued to control levels, while for flies fed 5 mM TMG, *Dm*OGT decreased below levels seen in the control genotype. To assay whether the same pharmacological rescue could be performed during development, adults were allowed to lay eggs on food supplemented with TMG and the O-GlcNAcylation levels of their offspring were measured by Western blot at the wandering third instar stage (Fig. 2C). Presumably due to differences in feeding behaviour, TMG concentrations required to rescue O-GlcNAcylation during the larval stages of development were much lower than for adults. At 150 μM TMG, O-GlcNAcylation in *sxc*^*C941Y*^ larvae was no longer significantly different from the control genotype, while O-GlcNAcylation in larvae fed 200 μM TMG was both significantly higher than in the *sxc*^*C941Y*^ larvae fed standard food and not significantly different from the control genotype. Overall, these results demonstrate that defective O-GlcNAc homeostasis in flies carrying an OGT-CDG mutation can be restored by reducing OGA activity through pharmacological inhibition.

### sxc^C941Y^ flies possess a neuromuscular junction bouton phenotype

Previous research has identified an important role for O-GlcNAcylation in excitatory synapse function^28, 47, 50^. To ascertain the contribution of this role of O-GlcNAcylation to ID, synaptic development was assayed at the larval neuromuscular junction (NMJ). This synapse is an established model for mammalian central nervous system (CNS) excitatory synapses and has been previously used to study the role of genes implicated in ID^52^. To assay the effects of *sxc* mutations on NMJ morphology, type 1b NMJs of muscle 4 were visualised by immunostaining for the subsynaptic reticulum protein Disks large 1 (Dlg1)^53^ and with an anti-HRP antibody to visualise neuronal membranes^54^ (Fig. 3A). Upon quantification with a semiautomated ImageJ macro^55^, several parameters measured were found to significantly differ between the NMJs in control genotype larvae and *sxc*^C941Y^ and the catalytically dead *sxc*^K872M^ larvae. The average NMJ area in *sxc^WT^* larvae (mean ± standard deviation, 326 ± 52 μm^2^) was significantly higher than in both *sxc*^*C941Y*^ (278 ± 28 μm^2^) and *sxc^K872M^* mutant larvae (198 ± 32 μm^2^), with a significant difference between the two *sxc* mutant groups. This phenotype was partially rescued in the *sxc*^*C941Y*^;*Oga^KO^* line (291 ± 39 μm^2^), relative to the control genotype, although the total area of the NMJs was not affected in the *Oga^KO^* larvae (337 ± 32 μm^2^), consistent with previous research on *Oga*^KO^ larvae ^47^ (Fig. 3B). Total length was also significantly different between the control genotype (mean ± standard deviation, 115 ± 18 μm) and *sxc*^*C941Y*^ (93 ± 11 μm) and *sxc^K872M^* larvae (74 ± 7 μm). This parameter was also partially rescued in *sxc*^*C941Y*^;*Oga^KO^* larvae (102 ± 14 μm) relative to *sxc^WT^* larvae, while being unaffected in the *Oga^KO^* genotype (115 ± 13 μm) (Fig. 3C). Finally, bouton numbers were also significantly reduced in both *sxc*^*C941Y*^ (mean ± standard deviation, 15 ± 3) and *sxc^K872M^* (12 ± 1) larvae, relative to the *sxc^WT^* controls (19 ± 3). Unlike total area and length, this parameter remained significantly reduced in the *sxc*^*C941Y*^;*Oga^KO^* line (16, ± 2) relative to the control genotype (Fig. 3D).

**Figure 3.**
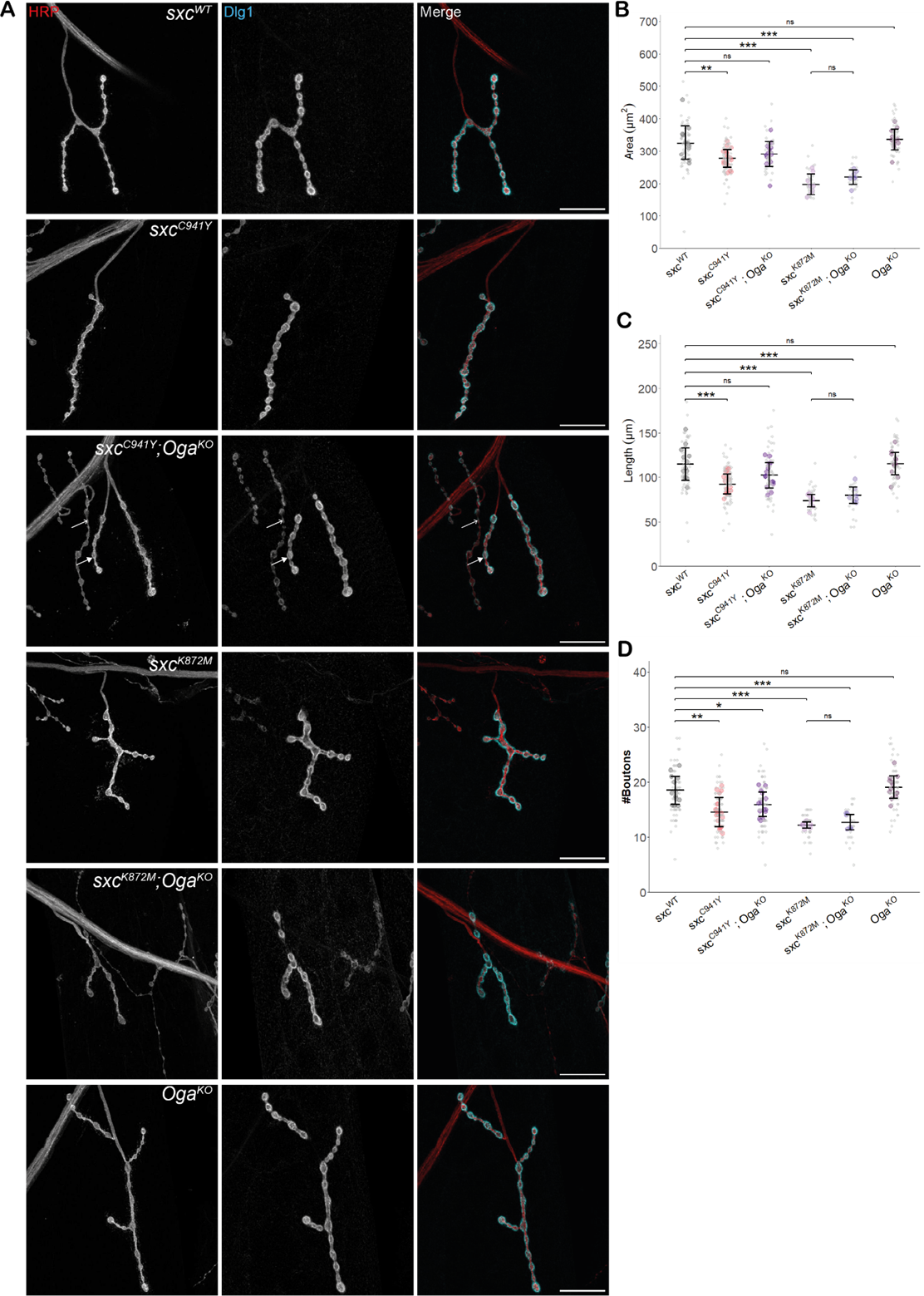
**A**) Representative images of larval neuromuscular junctions (NMJs) immunolabelled with anti-HRP (red), anti-Disks Large 1 (cyan) and both (scale bars 25 μm) for *sxc^WT^* (n = 13), *sxc*^*C941Y*^ (n = 19), *sxc*^*C941Y*^;*Oga^KO^* (n = 14), *sxc^K872M^* (n = 8), *sxc^K872M^;Oga^KO^* (n = 7) and *Oga^KO^* (n = 13) larvae. In the *sxc*^*C941Y*^;*Oga^KO^* panel the closed arrow indicates 1b boutons, analysed here, while the open arrow indicates an example of 1s boutons, not analysed here. **B)** Quantification of NMJ area (mean ± SD), which was found to be significantly different between genotypes (F(5,68) = 23.05, *p* < 0.001). Relative to the *sxc^WT^* control, both *sxc*^*C941Y*^ and *sxc^K872M^* larvae presented with a smaller NMJ area (*p_adj_* < 0.01 and *p_adj_* < 0.001, respectively), which was partially rescued in the *sxc*^*C941Y*^;*Oga^KO^* strain (*p_adj_* = 0.14), though the *Oga^KO^* larvae did not present with a significantly increased NMJ area (*p_adj_* = 0.97). **C)** Quantification of NMJ length (mean ± SD), which was found to be significantly different between genotypes (F(5,68) = 17.75, *p* < 0.001). Relative to the control genotype, both *sxc*^*C941Y*^ and *sxc^K872M^* larvae presented with overall shorter NMJ length (*p_adj_* < 0.001 for both), while NMJ length was not significantly different in *sxc*^*C941Y*^;*Oga^KO^* larvae (*p_adj_* = 0.13), despite *Oga^KO^* NMJ length not being affected (*p_adj_* = 0.99). **D)** Bouton number (mean ± SD) is significantly reduced in *sxc*^*C941Y*^ and *sxc^K872M^* larvae (F(5,68) = 18.11, p < 0.001, p_adj_ <0.001 for both), and remains significantly reduced in *sxc*^*C941Y*^;*Oga^KO^* larvae (*p_adj_* < 0.05). Values for individual NMJs are represented as small grey points, with averages for each larva represented as larger coloured points. Descriptive and inferential statistics were performed on larval averages, * p < 0.05, ** p < 0.01, *** p < 0.001.

As O-GlcNAcylation has been shown to regulate overall body size ^56, 57^ and NMJ area correlates with muscle size ^55^, we decided to measure muscle size in *sxc^K872M^* larvae to determine whether changes in overall body growth could explain the NMJ phenotype we observed. No significant difference in muscle size was observed between *sxc^WT^* and *sxc^K872M^* larvae, and when NMJ area was normalised to muscle area, this parameter remained significantly reduced in *sxc^K872M^* larvae (Fig. S3A). Overall, growth of larval neuromuscular junctions is broadly stunted in larvae modelling OGT-CDG and in larvae completely lacking OGT catalytic activity, with the phenotype partially rescued in the former by knocking out *Oga*. This is at odds with previously published research, which shows that both rationally designed hypomorphic mutants and ID mutations in the TPR domain result in increased growth at the neuromuscular junction ^47^. To address this disparity, we measured NMJ parameters in larvae of one of the genotypes previously assayed, *sxc^H596F^*. We found that this mutation also results in a significant decrease in NMJ area (mean ± standard deviation, 260 ± 23 μm^2^) relative to the control genotype (304 ± 18 μm^2^), with a similar effect for length and bouton number, consistent with the other genotypes assayed here (Fig. S3B-E).

### Pharmacological rescue of OGT-CDG neuromuscular junction phenotypes

To determine whether the (partial) rescue of NMJ parameters by genetic ablation of OGA activity can be recapitulated by pharmacological means, larvae were fed 200 μM TMG to elevate O-GlcNAcylation to control levels, as previously determined (Fig. 2C). As with knocking out *Oga*, elevating O-GlcNAcylation pharmacologically resulted in a partial rescue of NMJ parameters (Fig. 4A). The total NMJ area in *sxc*^*C941Y*^ larvae treated with 200 μM TMG (mean ± standard deviation, 303 ± 40 μm^2^) was no longer significantly different relative to the control genotype (319 ± 31 μm^2^) while *sxc*^*C941Y*^ fed a vehicle control presented with reduced NMJ area relative to the control genotype (272 ± 40 μm^2^) (Fig. 4B). Unlike in *sxc*^*C941Y*^;*Oga^KO^* larvae, TMG inhibition in *sxc*^*C941Y*^ larvae did not significantly rescue NMJ length (median ± interquartile range, 106 ± 13 μm) relative to the control genotype (117 ± 9 μm), although a non-significant increase in length relative to *sxc*^*C941Y*^ larvae fed a vehicle was observed (95 μm ± 8) (Fig. 4C). Similar to the OGA knockout experiment (Fig. 3D), *sxc*^*C941Y*^ larvae fed 200 μM TMG presented with significantly fewer boutons per NMJ (mean ± standard deviation, 16 ± 2) relative to the control genotype (19 ± 2) without a significant difference relative to the *sxc*^*C941Y*^ larvae fed a vehicle control (15 ± 2) (Fig. 4D). Overall, this demonstrates that pharmacological inhibition of OGA activity can partially rescue synaptogenesis in OGT-CDG mutant larvae.

**Figure 4.**
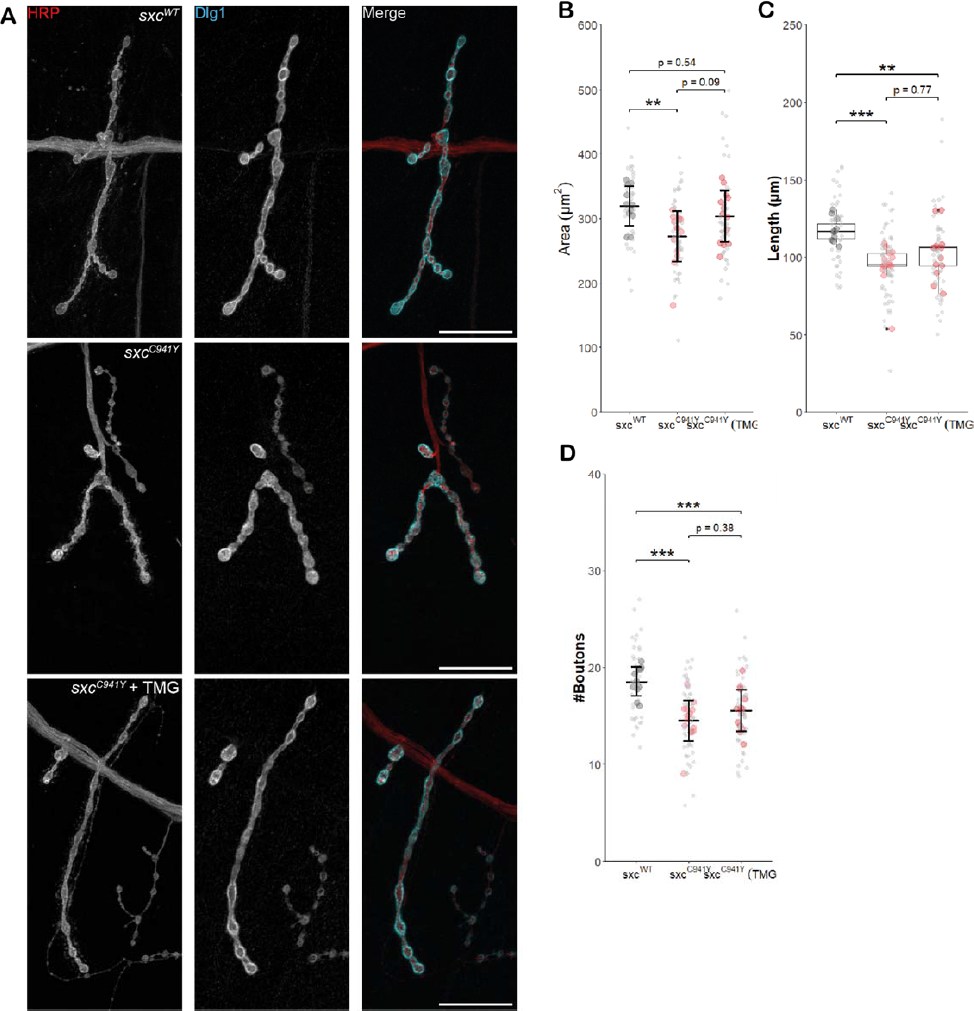
**A**) Representative images of NMJs immunolabelled with anti-HRP (red), anti-Disks Large 1 (cyan) and both (scale bars 25 μm) for *sxc^WT^* (n = 11), *sxc*^*C941Y*^ (n = 14), and *sxc*^*C941Y*^ fed 200 μM TMG (n = 13). **B**) NMJ area (mean ± SD) is significantly reduced in *sxc*^*C941Y*^ larvae fed a vehicle control (F(2,35) = 5.264, *p* = 0.01, *p_adj_* < 0.01), relative *sxc^WT^* larvae. s*xc*^*C941Y*^ larvae fed 200 μM TMG no longer present with a significant reduction in total NMJ area (*p_adj_* = 0.54). **C)** Total NMJ length (median ± IQR) is significantly different between groups (Χ^2^ (2) = 17.483, p < 0.001), however, unlike total area, post hoc analysis demonstrates that this parameter remains significantly reduced compared to the *sxc^WT^* control for both vehicle and TMG treated *sxc*^*C941Y*^ larvae (p_adj_ < 0.001 and p_adj_ < 0.01, respectively). **D**) Quantification of bouton number (mean ± SD) demonstrated a significant intergroup difference (F(2,35) = 13.6, *p* < 0.001), with both vehicle and TMG fed *sxc*^*C941Y*^ larvae presenting with significantly reduced bouton number (*p_adj_*< 0.001 and *p_adj_* < 0.01, respectively). Values for individual NMJs represented as small grey points, with averages for each larva represented as larger coloured points. Descriptive and inferential statistics were performed on larval averages, * p < 0.05, ** p < 0.01, *** p < 0.001.

### Fragmented sleep in sxc^C941Y^ flies is reversable by normalising global O-GlcNAcylation

Patients with ID present with hyper-activity and sleep disturbances more often than the general population^58, 59^. Several patients affected by OGT-CDG follow this pattern, presenting with sleep disturbances and behavioural abnormalities^4, 8^. To assay whether activity and sleep are also disrupted in a *Drosophila* model of OGT-CDG, we used the *Drosophila* Activity Monitor (DAM) to measure these parameters (Fig. 5A,B). In *Drosophila* research, sleep is commonly defined as a period of five or more minutes of quiescence, which is accurately measured by the DAM system^60^. Total activity of *sxc*^*C941Y*^ flies (median ± interquartile range, 1.25e3 ± 6.6e2 counts/24 h) was not significantly different from the control genotype (1.23e3 ± 5.8e2 counts/24 h). However, *sxc*^*C941Y*^;*Oga^KO^* flies were significantly less active than the control genotype (8.9e2 ± 3.9e2 counts/24 h), despite the *Oga^KO^* allele having no effect on total activity on its own (1.13e3 ± 5.7e2 counts/24 h) (Fig. 5C). By contrast, *sxc*^*C941Y*^ flies did present with reduced total sleep (mean ± standard deviation, 8.1e2 ± 1.8e2 min/24 h), relative to the control genotype (9.4e2 ± 1.4e2 min/24 h), which was rescued in *sxc*^*C941Y*^;*Oga^KO^* flies to wild type levels (9.7e2 ± 1.3e2 min/24 h) (Fig. 5D). Upon more detailed investigation, the nature of sleep disruption in the OGT-CDG flies was found to be due to a reduced duration of individual sleep bouts in these flies (median ± interquartile range, 32 ± 15 min) compared to the control genotype (53 ± 30 min). Mean sleep bout duration in *sxc*^*C941Y*^ flies is partially rescued by elevating global O-GlcNAcylation through knocking out *Oga* (40 ± 23 min), although it remained significantly reduced compared to the control genotype (Fig. 5E). Further investigation of sleep bout duration, we found that the differences in sleep patterns between genotypes could be explained by the inability of *sxc*^*C941Y*^ flies to maintain longer sleep bouts. *sxc^WT^* flies experience significantly more sleep bouts longer than 2 h (median ± interquartile range 2.0 ± 1.0 bouts/24 h) relative to *sxc*^*C941Y*^ flies (1 ± 1.3 bouts). This aspect of sleep is also rescued by knocking out *Oga*, with *sxc*^*C941Y*^;*Oga^KO^* flies no longer presenting with a significant decrease in number of sleep bouts longer than 2 h (1.7 ± 1.3 bouts/24 h) (Fig. 5G). Accompanying decreased sleep bout duration, *sxc*^*C941Y*^ and *sxc*^*C941Y*^;*Oga^KO^* flies present with significantly more frequent sleep bouts (27 ± 7 and 26 ± 8 bouts/24 h, respectively) than the control genotype (19 ± 7 bouts/24 h) (Fig. 5F). These results indicate that the sleep defects in *sxc*^*C941Y*^ flies are only partially rescued by elevating global O-GlcNAcylation, with the modest rescue of sleep bout duration seen upon loss of OGA fully rescuing total sleep, in part due to sleep frequency remaining unaltered and above the control genotype levels.

**Figure 5.**
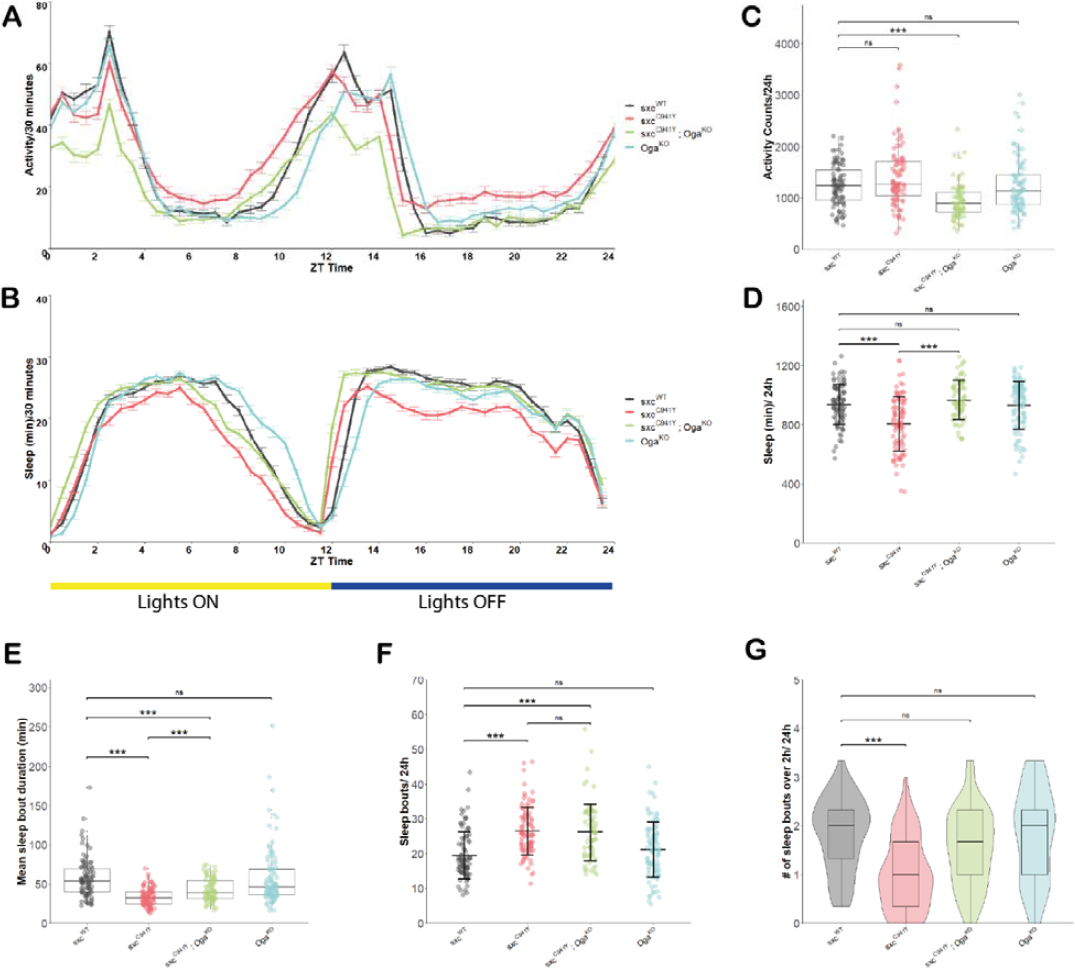
**A**) Activity profile (mean ± SEM of activity counts in 30 minute bins) for *sxc^WT^* (n = 89), *sxc*^*C941Y*^ (n = 94), *sxc*^*C941Y*^;*Oga^KO^* (n = 74), and *Oga^KO^* (n = 95) flies. Sleep profile (mean ± SEM of sleep in 30 minute bins) for genotypes as in **A**. **C-G)** Sleep parameters for genotypes in **A** and **B. C)** Total daily activity (median ± IQR) is significantly reduced in the *sxc*^*C941Y*^;*Oga^KO^* mutant strain relative to the control (Χ^2^ (3) = 41.546, *p* < 0.001, *p_adj_* <0.001). **D)** Total daily sleep (mean ± SD) is significantly reduced in *sxc*^*C941Y*^ flies relative to the control genotype (F(3,348) = 18.34, *p* < 0.001, *p_adj_* < 0.001), while both *sxc*^*C941Y*^;*Oga^KO^* and *Oga^KO^* flies do not have significantly altered total sleep (*p_adj_* = 0.58 and *p_adj_*= 0.99, respectively). **E)** Mean sleep episode duration (median ± IQR) is significantly reduced in both *sxc*^*C941Y*^ and *sxc*^*C941Y*^;*Oga^KO^* flies relative to the control genotype (Χ^2^ (3) = 75.084, *p* < 0.001, *p_adj_* < 0.001 for both). Mean sleep episode duration is significantly increased in *sxc*^*C941Y*^;*Oga^KO^* flies compared to *sxc*^*C941Y*^ flies (*p_adj_* < 0.001). **F)** Daily number of sleep bouts (mean ± SD) is significantly elevated in both *sxc*^*C941Y*^ and *sxc*^*C941Y*^;*Oga^KO^* flies compared to the *sxc^WT^* control (F(3,348) = 20.31 *p* < 0.001, *p_adj_* < 0.001 for both)**. G)** Daily number of sleep bouts longer than 2 h (median ± IQR) is significantly lower in *sxc*^*C941Y*^ flies than the control genotype (Χ^2^ (3) = 49.623, *p* < 0.001, *p_adj_* < 0.001). Individual points represent mean values of measurements conducted over three days, for unique flies. * p < 0.05, ** p < 0.01, *** p < 0.001.

To dissect developmental from non-developmental contributions to this sleep phenotype, we investigated whether elevating O-GlcNAcylation only in adulthood could rescue the sleep phenotype observed in *sxc*^*C941Y*^ flies. Adult *sxc*^*C941Y*^ flies were fed 3 mM TMG for 72 h prior to and during activity monitoring. In this condition, OGT-CDG flies no longer presented with decreased overall sleep duration (Fig. 6A). However, other aspects of sleep remained disrupted in OGT-CDG flies. Both mean sleep duration (median ± interquartile range, 31 ± 12 min) and daily number of sleep bouts longer than 2 h (median ± interquartile range, 1 ± 1 bouts/24 h) remained significantly reduced compared to the *sxc^WT^* control (37 ± 14 min, 1.3 ± 1.0 bouts/24h, respectively). Additionally, as in previous experiments, *sxc*^*C941Y*^ flies presented with significantly more sleep bouts (mean ± standard deviation, 32 ± 8 bouts/24 h) than the control genotype (26 ± 6 bouts/24 h) (Fig 6B-D). Interestingly, these phenotypes were partially reversed by TMG feeding. Mean sleep bout duration in *sxc*^*C941Y*^ flies fed TMG was no longer significantly different from the control genotype (34 ± 14 min), nor was the number of sleep bouts longer than 2 h (1.3 ± 1.0 bouts/24 h). The number of sleep bouts in *sxc*^*C941Y*^ flies fed TMG was significantly fewer than in the same genotype fed a vehicle control (29 ± 7 bouts/24 h), although it remained non-significantly elevated relative to the control genotype. These results suggest that effects of OGT-CDG mutations may not be solely developmental, and that defective O-GlcNAc cycling in adulthood may be an important contributor to the pathogenesis of these mutations.

**Figure 6.**
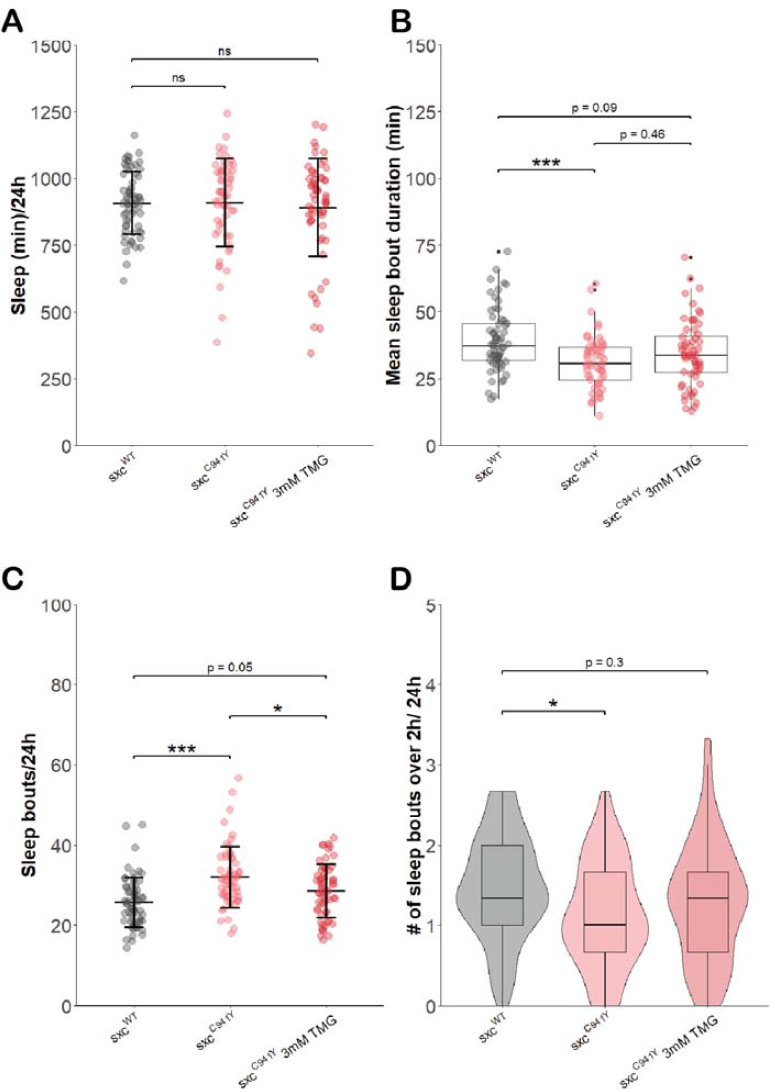
Sleep parameters for *sxc^WT^* (n = 63), *sxc*^*C941Y*^ (n = 57) and *sxc*^*C941Y*^ flies fed 3 mM TMG (n = 60). **A**) Total sleep (mean ± SD) is not significantly different between groups (F(2,177) = 0.249, *p* = 0.78). **B)** Mean sleep bout duration (median ± IQR) significantly differs between groups (Χ^2^ (2) = 49.623, *p* < 0.001), though is only significantly reduced for *sxc*^*C941Y*^ flies fed the vehicle control (*p_adj_* <0.001) and not TMG (*p_adj_* = 0.09). **C)** Daily number of sleep bouts is significantly increased in *sxc*^*C941Y*^ flies fed a vehicle control and is significantly reduced by TMG feeding (F(2,177) = 13.11, *p* < 0.001, *p_adj_* < 0.001 and *p_adj_* < 0.05, respectively). **D)** The number of sleep bouts longer than 2 h is significantly reduced in *sxc*^*C941Y*^ flies fed a vehicle control, but not TMG (Χ^2^ (2) = 8.2491, *p* < 0.05, *p_adj_* <0.05 and *p_adj_* = 0.3, respectively). Individual points represent mean values of measurements conducted over three days, for unique flies. * p < 0.05, ** p < 0.01, *** p < 0.001.

## Discussion

Many mutations in *OGT* causal in intellectual disability modelled previously do not result in a decrease in global O-GlcNAcylation either in embryonic stem cells or patient derived fibroblasts, in many cases due to feedback mechanisms reducing OGA protein levels^3, 5, 8^. The mutation modelled here is one of only two which has been shown to reduce global O-GlcNAcylation when modelled in mammalian cells^6, 22^. Here, we have shown that patient mutations in the catalytic domain of OGT result in decreased O-GlcNAcylation in adult flies, corroborating previous results^5^. Expanding upon these results, we have demonstrated that an OGT-CDG catalytic domain mutation can reduce O-GlcNAcylation throughout development, despite a compensatory increase in total *Dm*OGT protein. When modelled in mouse embryonic stem cells, this mutation (C921Y in humans and mice) also results in an increase in OGT protein levels^22^. It is tempting to assert that increased OGT, as opposed to decreased OGA, is a homeostatic mechanism linked specifically to this mutation. However, in the fly, increased *Dm*OGT protein levels appear to be a response commonly associated with decreased OGT catalytic activity, demonstrated here by the *sxc^K872M^* mutant stain and in previous work^5^. While this increase in *Dm*OGT protein levels was not explored in further detail, some evidence exists for post-transcriptional regulation of *sxc* expression through alternate splicing^61^, a mechanism known be involved in the control of *OGT* and *OGA* expression and O-GlcNAc homeostasis in mammalian cells^62^. We have also shown that reduced catalytic activity of *Dm*OGT as a result of modelling a patient mutation in *sxc* can phenocopy rational mutagenesis of a key catalytic residue (*sxc^H537A^*) causing the growth of ectopic scutellar bristles^48^. Also known as macrochaetae, the development of these sensory cells is well studied, particularly in the context of cell fate determination by lateral inhibition through Notch signalling^63^, providing a tractable system for the understanding of the impacts of hypo-O-GlcNAcylation on cell fate determination. With reports of Notch signalling requiring appropriate O-GlcNAcylation^30^, this phenotype presents an interesting system to research the contribution of the Notch signalling pathway to ID.

A key question regarding OGT-CDG is whether therapeutic approaches targeting OGA can raise O-GlcNAcylation and potentially ameliorate symptoms in this disorder, as previously proposed^4^. Here, we show that normal global O-GlcNAcylation levels can be restored in *sxc*^*C941Y*^ adult flies through knockout out of *Oga.* Previous research has demonstrated that *Oga^KO^* alleles can rescue phenotypes associated with reduced O-GlcNAcylation^47^, however, here we present the first direct evidence that global O-GlcNAcylation can exceed control levels in *Dm*OGT hypomorphic flies in the absence of OGA. While we did not see a concomitant rescue of the scutellar bristle phenotype seen in *sxc*^*C941Y*^ flies, this may occur as a consequence of unique kinetics of O-GlcNAcylation and removal of O-GlcNAc on various *Dm*OGT substrates – i.e., the dysregulation of the ratio of stoichiometries of modification of specific substrates may be exacerbated in the absence of OGA. Previous research has demonstrated this may occur in mammalian cells, with some O-GlcNAcylated proteins not affected by OGA inhibition in cancer cells^64, 65^. Alternatively, it may be that both the addition and timely removal of O-GlcNAc from specific proteins is required for normal scutellar bristle development.

Complete ablation of *Oga* expression is a blunt approach, elevating O-GlcNAcylation levels beyond those seen in the control genotype and is not a feasible therapeutic approach. Pharmacological approaches to inhibit OGA offer more precise control over the degree of O-GlcNAcase activity and are being actively pursued as potential treatments for neurodegenerative disorders^66^. Our experiments suggest that the potent OGA inhibitor TMG can be used to rescue global O-GlcNAcylation levels in *sxc*^*C941Y*^ flies to those of a genetic background control, at various stages of development. Interestingly, rescuing O-GlcNAcylation levels through OGA inhibition also restored OGT levels in *sxc*^*C941Y*^ flies. However, there is a clear difference in the pattern of O-GlcNAcylation visualised by immunoblotting in adult *sxc*^*C941Y*^ flies fed TMG relative to the control genotype. This incomplete rescue of O-GlcNAcylation may be consequential in phenotypic rescue. Additionally, a paradoxical effect was seen upon feeding higher doses of TMG. Because inhibition of OGA appears to reduce protein levels of OGT, global O-GlcNAc levels were not rescued at higher levels of TMG. However, specific immunoreactive bands appeared to maintain elevated levels of O-GlcNAc. This potentially indicates that mechanisms controlling OGT expression in response to O-GlcNAcylation levels are particularly sensitive to OGA activity, lowering OGT protein levels prior to O-GlcNAcylation stoichiometry rising on some OGT substrates.

Previous research has shown that alleles encoding hypomorphic variants of *Dm*OGT result in overgrowth at the neuromuscular junction, and that TPR domain mutations modelling those seen in patients result in a similar phenotype^47^. However, here, we observe the opposite effect, with both *sxc*^*C941Y*^ and *sxc^K872M^* larvae presenting with smaller NMJs, likely explained by impaired addition of boutons. This discrepancy is unlikely to be caused by allele specific effects as this hypothesis was tested by assaying NMJ parameters in one of the hypomorphic mutants previously described and finding that this mutation (*sxc^H596F^*) also results in stunted growth at the NMJ. While we utilised different markers to count boutons than in previous research^47^, discrepancies in area and length of NMJs cannot be explained in this manner. Nonetheless, as in previous research on the effects of reduced OGT catalytic activity on NMJ morphology, knocking out *Oga* can partially rescue phenotypes at this type of synapse^47^. This phenotype can also be partially rescued through pharmacological means, through the use of TMG. This could serve as proof of principle that pharmacological intervention in OGT-CDG is possible. This is not an immediately obvious conclusion, as it is possible that beyond the stoichiometry of the modification on individual substrates, the timing of addition and removal of O-GlcNAcylation could be important for synaptic development and function.

We also demonstrate a novel behavioural effect resulting from catalytic domain mutations in *sxc*. Normal O-GlcNAc cycling is required for maintenance of sleep in adult flies, and reduced *Dm*OGT catalytic activity as a result of an OGT-CDG mutation results in shorter, more frequent sleep episodes. Previously, a patient with this condition was reported to suffer from sleep disturbances characterised by abnormal EEG during sleep and insomnia^8^. This is particularly relevant given that normal sleep is required for multiple cognitive processes, such as memory formation, both in flies and humans^67–69^. Encouragingly, this phenotype can be partially rescued by knocking out OGA or by pharmacologically elevating O-GlcNAcylation in adult flies. This result could have important implications for our understanding of OGT-CDG, providing the first evidence that suggests that the disorder is not purely developmental and may be amenable to therapeutic approaches at later stages of life.

## Methods and Materials

### CRIPSR-Cas9 mutagenesis

The gRNA sequence for generating the sxc C941Y flies was selected using the online tool Crispr.mit.edu. The optimal gRNA sequence was included in annealing oligos including overhangs compatible with cloning into the pCFD3-dU63gRNA plasmid previously cut with BpiI restriction enzyme. A 2kb repair template for the region was generated from Drosophila Schneider 2 cell genomic DNA by PCR using GoTaq G2 Polymerase. The PCR product was cloned as a blunt product into the pTOPO-Blunt plasmid. Mutations were introduced into the template to include the C941Y mutation as well as silent mutations to remove the gRNA recognition sequence. This was carried out using the QuikChange kit from Stratagene and confirmed by DNA sequencing. The mutations removed the restriction site BseMI which is present in the gRNA sequence*. sxc*^C941Y^ mutant flies were generated by microinjection of vas-Cas9 embryos (BL51323) (Rainbow Transgenic Flies, Inc) with CRISPR reagents generated in-house, backcrossed to a w^1118^ (VDRC60000) background and the mutated chromosome was balanced over Curly of Oster (CyO). Diagnostic digests were carried out on the resulting flies to first confirm the loss of the restriction site followed by sequencing of the PCR product. The correctness of the mutation was also confirmed through sequencing of the full-length sxc mRNA.

### Fly stocks and maintenance

Stocks were maintained on a 12:12 light dark cycle at 25 °C on Nutri-Fly Bloomington Formulation fly food. *sxc^K872M^* mutant flies from Mariappa et al. (2018) were used. *Oga^KO^* flies from Muha et al. (2020) were used to generate *sxc*^*C941Y*^;*Oga*^*KO*^ and *sxc^K872M^;Oga^KO^* stocks. The homozygous lethal *sxc^K872M^* chromosome was balanced over a CyO chromosome carrying a GFP reporter (CyO, P{ActGFP.w[-]}CC2, BL9325). An isogenic w^1118^ (VRDC60000) background strain was used as a control genetic background.

### Drosophila tissue lysis and Western blotting

For immunoblotting of adult head lysates, flies raised as described previously were anaesthetised with CO_2_ and an equal number of 3–5-day old male and female flies were snap frozen in liquid nitrogen. Heads were then severed from bodies by vortexing flies twice and collected using a paintbrush. To collect larval and embryonic lysates, homozygous 3-5 day old females and males were allowed to lay embryos for 4 h and 2 h, respectively, on apple juice agar plates supplemented with yeast paste. For recessive lethal lines, heterozygous parents were crossed in the same manner. Embryos were collected 14 h later and snap frozen on dry ice. For recessive lethal genotypes, homozygous embryos were collected based on the absence of a GFP fluorescent CyO balancer chromosome. For larval tissues, 24 h after embryo collection, *sxc* mutant homozygous first instar larvae were collected into vials containing Nutri-Fly Bloomington Formulation fly food at a density of 25 larvae per vial and aged to the wandering third instar stage, when they were snap frozen on dry ice. For experiments in which specificity of the O-GlcNAc antibody was tested by prior incubation with *Clostridium perfringens OGA CpOGA,* heads were lysed in modified RIPA buffer to accommodate the pH optimum of *Cp*OGA ^70^ (150 mM NaCl, 1% NP-40, 0.5% sodium deoxycholate, 0.1% SDS, 25 mM citric acid pH 5.5) supplemented with a protease inhibitor cocktail (1 M benzamidine, 0.2 mM PMSF, 5 mM leupeptin). To validate specificity of O-GlcNAc detection, lysates were split with one group incubated with 2.5 μM GST tagged *cp*OGA to remove O-GlcNAc while the experimental group was incubated with 1 μM GlcNAcstatin G. Lysates were then incubated for 2 h at room temperature, agitated at 300 RPM using a thermomixer (Eppendorf thermomixer comfort). The reaction was stopped by heating to 95 °C with NuPAGE LDS Sample Buffer with 50 mM TCEP to a 1x concentration. Otherwise, collected tissues were lysed in 50 mM Tris-HCl (pH 8.0), 150 mM NaCl, 1 % Triton-X 100, 4 mM sodium pyrophosphate, 5 mM NaF, 2 mM sodium orthovanadate, 1 mM EDTA, supplemented 1:100 with a protease inhibitor cocktail (1 M benzamidine, 0.2 mM PMSF, 5 mM leupeptin) and 1.5x NuPAGE LDS Sample Buffer with 50 mM TCEP. Protein concentration was estimated using a Pierce 660 assay (Thermo Scientific) supplemented with ionic detergent compatibility reagent (Thermo Scientific). 30 μg of protein per group were separated by gel electrophoresis (NuPage 4-12% Bis-Tris, Invitrogen) and transferred onto a nitrocellulose membrane (Amersham Protran 0.2 μm). Membranes were developed with the following primary antibodies: mouse anti-O-GlcNAc (RL2, 1:1000, Novus), rabbit anti-OGT (1:1000, Abcam, ab-96718) and rabbit anti-actin (1:5000, Sigma, A2066) and the following secondary antibodies: goat anti-mouse IgG 800 and donkey anti-rabbit IgG 680 infrared dye conjugated secondary antibodies (Li-Cor, 1: 10,000). Western blots were analysed using Image Studio Lite.

### Thiamet G feeding

Thiamet G (SantaCruz, sc-224307) was dissolved in PBS to a stock concentration of 100 mM. This stock was mixed with *Drosophila* instant food (Flystuff Nutri-Fly Food, Instant Formulation) to appropriate concentrations, to avoid heating Thiamet G. For experiments with adult flies, 1-3 day old flies (males and females in equal proportion) were placed on food for 72 h prior to snap freezing in liquid nitrogen. For larval feeding experiments, 10 0-3 day old females were crossed with 4 males and allowed to lay embryos for 2 days. Wandering 3^rd^ instar larvae were snap frozen on dry ice and lysed.

### Neuromuscular Junction Immunohistochemistry

The neuromuscular junction (NMJ) assay was performed as in Nijhof et al. (2016). Larvae for this assay were obtained as described above. Male wandering third instar larvae were dissected using the ‘open book’ technique ^71^ followed by immediate fixation in 3.7% paraformaldehyde in phosphate buffered saline (pH 7.5) (PBS) for 25 min. Fixed larvae were either stored in PBS at 4 °C for up to 48 h or immediately processed further. Larval preparations were blocked using 5% normal donkey serum (NDS) in PBS and Triton-X (0.3%, PBST) for 2 h at room temperature, followed by immunostaining using: mouse anti-Disks Large 1 (1:25, Developmental Studies Hybridoma Bank, RRID: AB_528203) and goat anti-HRP conjugated to Alexa Fluor 647 (1:400, Jackson ImmunoResearch, RRID: AB_2338967) in 5% NDS PBST overnight at 4 °C. Sections were washed 4 times for 10 min in PBST (0.5%), followed by 4 h incubation with donkey anti-mouse Alexa Fluor 488 in 5% NDS PBST at room temperature. Sections were washed as for primary antibodies, rinsed in PBS, and mounted using Dako Fluorescence Mounting Medium (Agilent). Images of type 1b NMJs of muscle 4 were obtained using a Zeiss 710 confocal microscope using a 10x objective (EC Plan Neofluar 0.3) (voxel size: 0.69 x 0.69 x 6.22 μm) for muscle area measurements and using a 63x objective (Plan Apochromat 1.4 oil) to image individual junctions (voxel size: 0.196 x 0.196 x 0.91 μm). Image size for the former was 2048×2048 pixels and 688×688 for the latter. Both channels were acquired simultaneously. NMJ parameters were scored using a semi-automated macro (Neuromuscular Junction Morphometrics ^55^) with poorly annotated or damaged NMJs excluded from further analysis. Muscle area was manually measured using the polygon selection tool in ImageJ. Statistical analysis was performed on mean values for individual larvae for which 3 or more NMJs were accurately annotated.

### Drosophila activity monitor

*Drosophila* activity was recorded using Trikinetics DAM2 monitors. 1–3-day old male flies were used for all experiments. Briefly, male flies were anaesthetised using CO_2_ and placed in DAM vials with Nutri-Fly Bloomington Formulation food. Experiments were performed at 25°C on a 12 h:12 h light:dark cycle, data were recorded for 3 days, after 2 days of acclimatisation. For TMG rescue experiments, food was prepared as described in *Thiamet G feeding* and data were recorded 72 h after placing flies on supplemented food. Data were pre-processed using DAMFileScan113 software and Sleep and Circadian Analysis MATLAB Program (S.C.A.M.P.) ^60^. Data from at least 3 independent experiments were pooled for analysis. For analysis of number of bouts longer than 2 h the raw output from the DAM system was analysed in R using the Rethomics packages ^72^.

### Scutellar bristle assay

To assay scutellar bristle number, 8-10 young homozygous virgin females were mated with 3 males and allowed to lay embryos for 3 days to prevent overcrowding of larvae. Eclosed offspring were immobilised using CO_2_ and scutellar bristles were counted using a Motic SMZ-161 microscope.

### Western blot intensity profile

Intensity of O-GlcNAc immunoreactivity was calibrated to estimated molecular weight plotted using custom Python code (available at https://github.com/IggyCz/WBplotProfile). Briefly, images were imported using the PIL library ^73^, converted to NumPy arrays ^74^ and molecular weight markers were identified as intensity peaks in a user defined x-coordinate column of pixels. The SciPy ^75^ library was then used to fit a curve to identified molecular weight markers, to infer molecular weights at y-pixel coordinates. This was then used to calibrate the x-axis for plotting (using the matplotlib library ^76^) the relative intensity of immunolabelling across genotypes and conditions based on user defined x pixel coordinates defining protein lanes, normalised to loading controls.

### Statistical analyses

All statistical analyses were performed in R (version 4.0.3). Data that satisfied assumptions regarding homoscedasticity and normality were analysed with a one-way ANOVA followed by Tukey’s HSD with Bonferroni correction. Otherwise, data were analysed using a Kruskal-Wallis rank sum test followed by pairwise comparisons using Wilcoxon rank sum test with continuity correction and p value adjustment using the Bonferroni method. One outlier was removed from analysis (*sxc*^*N595K*^ OGT and O-GlcNAcylation quantification), based on the criteria of falling more than 1.5 interquartile range beyond the 75 percentile. To balance the removal of this outlier, the minimum for this group was also removed.

## Acknowledgements

This work was funded by a Wellcome Trust Investigator Award (110061) and a Novo Nordisk Foundation Laureate award (NNF21OC0065969) to D.M.F.v.A and a PhD studentship from the National Centre for the Replacement, Refinement and Reduction of Animals in Research (NC3Rs, award number T001682). We also thank Leeanne McGurk and Jens Januschke for their feedback as well as past and current members of our laboratory for their input, including Hannah Smith, Marta Murray, Veronica Pravata, and Conor Mitchell.

---

**Figure S1.**
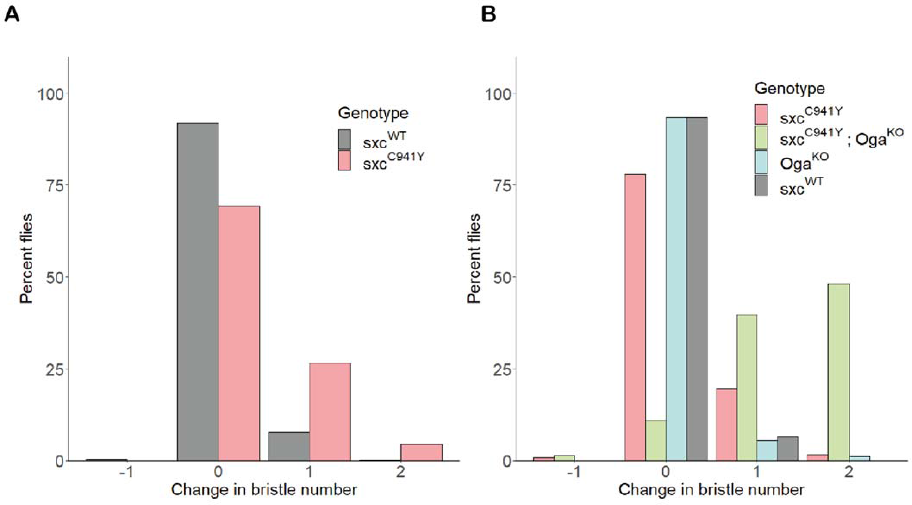
**A**) Quantification of the number of bristles on the scutellum of *sxc^WT^* (n = 566) and *sxc*^*C941Y*^ (n = 344) flies, represented as a percentage of total flies included in quantification. **B**) As for **A**, comparing the number of bristles for *sxc^WT^* (n = 136), *sxc*^*C941Y*^ (n = 123), *xc*^*C941Y*^;*Oga*^*KO*^ (n = 83) and *Oga^KO^* flies (n = 92).

**Figure S2.**
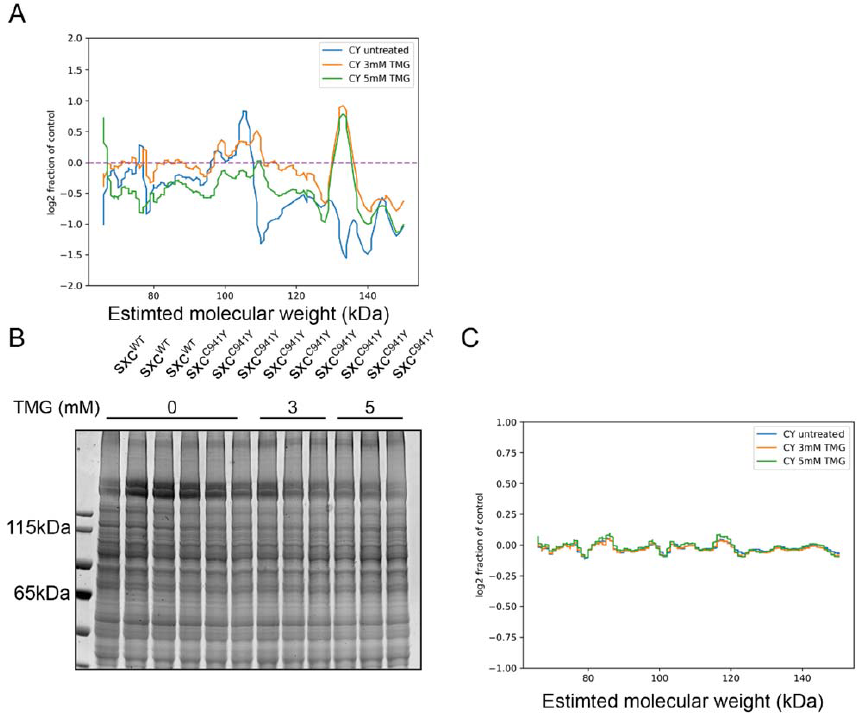
**A**) Log2 fraction of linear profile of O-GlcNAc immunoreactivity from the Western blot in Fig. 2B of *sxc*^*C941Y*^ vehicle (CY untreated), 3 mM Thiamet G (TMG) (CY 3 mM TMG) and 5 mM TMG (CY 5 mM TMG) relative to the *sxc^WT^* control (normalised to a loading control). Generated using a custom Python script to calibrate molecular weights to a curve fitted to the protein ladder. **B**) Coomassie stain of lysates used for the Western blot in Fig. 2B, and linear profile as in **A** (**C**).

**Figure S3.**
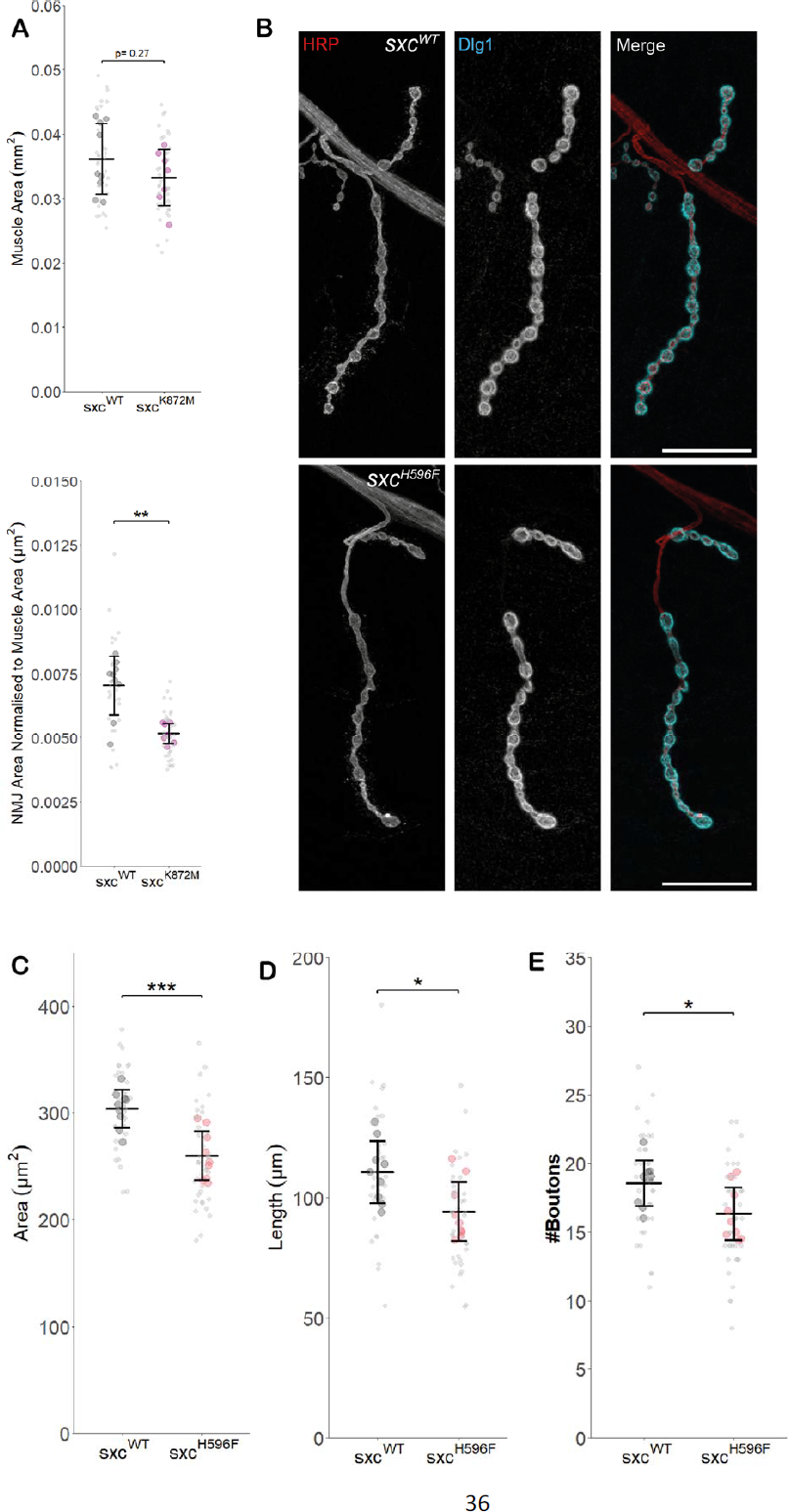
**A**) Muscle area from *sxc^WT^ (*n = 9, mean ± standard deviation 0.036 ± 0.005 mm^2^) and *sxc^K872M^* (n = 7, 0.033 ± 0.004 mm^2^, F(1,14) = 1.297, *p* = 0.27) larvae is not significantly different. When normalised to muscle area, NMJ area in *sxc^K872M^* larvae (0.0052 ± 0.0004 µm^2^ /µm^2^) is significantly reduced compared to *sxc^WT^* larvae (0.0071 ± 0.0012 µm^2^ /µm^2^, F(1,14) = 16.82, *p* < 0.01). **B**) Representative images of larval neuromuscular junctions (NMJs) immunolabelled with anti-HRP (red), anti-Disks Large 1 (cyan) and both (scale bars 25 μm) for *sxc^WT^* and *sxc^H596F^* larvae. **B-D**) Quantification of NMJ parameters quantified using a semi-automated ImageJ plugin for *sxc^WT^* (n = 9) and *sxc^H596F^* (n = 9) larvae, area (*sxc^WT^*:304 ± 18 µm^2^, *sxc^H596F^*: 260 ± 23 µm^2^, F(1,16) = 20.54, *p* < 0.001) (**B**), length (*sxc^WT^*:111 ± 12.8 µm, *sxc^H596F^*: 94.2 ± 12.3 µm, F(1,16) = 7.694, *p* < 0.05) (**C**), and bouton number (*sxc^WT^*: 18.5 ± 1.7, *sxc^H596F^*: 16.3 ± 1.9, F(1,16) = 6.864, *p* < 0.05) (**D**) are all significantly different between the two genotypes. Values for individual NMJs are represented by small grey points, with averages for each larva represented as larger coloured points. Descriptive and inferential statistics were performed on larval averages. * p < 0.05, ** p < 0.01, *** p < 0.001.

